# Increased Epstein-Barr virus C-promoter activity with CTCF-binding site deletion is associated with elevated EBNA2 recruitment

**DOI:** 10.1101/442186

**Authors:** Ian J Groves, Martin J Allday

## Abstract

The regulation of transcription from Epstein-Barr virus promoters is known to involve the association of the host CCCTC-binding factor (CTCF) protein. This control involves direct binding of CTCF across the EBV genome and the formation of three-dimensional loops between virus promoters and enhancers. We sought to address how the deletion of a CTCF binding site upstream of the C-promoter (Cp) affected viral transcription in infected lymphoblastoid cell lines (LCLs) and how binding of the EBV trans-activating protein EBNA2 was changed across this promoter. Transcript level from Cp was up-regulated with CTCF binding site deletion, and transcription from other promoters (Wp and Qp) was decreased, while transcript levels were largely unchanged by independent mutation of a Cp-RBPJκ binding site. In turn, expression of EBNA2 protein was also increased, likely driven by increases in polycistronic EBNA2-encoding transcripts. Finally, Cp up-regulation was associated with an 8-fold increase in EBNA2 enrichment across Cp, concomitant with increased association of the associated cellular factor RBPJκ, probably due to a more accessible three-dimensional chromatin conformation upstream of Cp. Overall, the data presented here confirm that binding of CTCF directly upstream of Cp is important for the regulation of transcription from this and other EBV promoters.

## Introduction

Epstein-Barr virus (EBV) is a member of the human gammaherpesvirus family and is commonly known for its association with infectious mononucleosis/glandular fever. However, EBV is also closely linked with as many as 1.5% of all human cancers including B-cell lymphomas and epithelial malignancies, such as Burkitt’s (BL) and Hodgkin’s lymphomas (HL) and nasopharyngeal carcinoma (NPC), respectively (Plummer, 2016; Farrell, 2018). The association of EBV with different diseases is characterised by distinct transcription profiles of EBV nuclear antigens (EBNAs) and latent membrane proteins (LMPs) in conjunction with the differentiation status of the infected B-cell. Latency type I, mostly associated with BL and proliferating infected memory B-cells, involves transcription from the Q-promoter (Qp) of EBNA1, a DNA binding protein that is able to tether the viral episome to cellular chromosomes, thereby maintaining the EBV genome in dividing cells (Westhoff Smith, 2013). In contrast, latency type II is associated with expression of LMP1 and LMP2 as well as Qp-driven EBNA1. Finally, latency type III (also known as the ‘growth’ program) comprises expression of the LMPs as well as EBNA1, 2, 3A-C and -LP via alternatively spliced polycistronic transcripts driven initially from the W-promoter (Wp), and subsequently from the C-promoter (Cp) (Rowe, 1987), partly due to EBNA2 recruitment (Woisetschlaeger, 1991; Altmann, 2006).

Programs of transcription during latency are driven from separate viral promoters on the EBV genome and are known to be regulated, at least in part, through modification of chromatin on the viral genome (Tempera, 2014; Hammerschmidt, 2015). The activity of Cp is also controlled by interaction with the ‘plasmid origin of replication’ (OriP), which acts as an enhancer (Reisman, 1986; Altmann, 2006; Puglielli, 2006). EBNA2 association further allows the recruitment and stimulation of RNA polymerase II (RNAPII) at Cp (Bark-Jones, 2006; Palermo, 2008). Recruitment of EBNA2 to Cp is thought to be, at least in part, directed by the presence of a binding site for RBPJκ, a host protein that binds to EBNA2 and co-localises with most EBNA2 binding sites on the genome (Ling, 1993; Zhao, 2011).

Interestingly, a host protein known as CCCTC-binding factor (CTCF) has been shown to be important in the regulation of gene expression of a number of human DNA viruses (Pentland, 2015) and is also able to bind to several locations on the EBV episome, including a latency III-specific site between OriP and Cp (Tempera, 2010). CTCF, a host genomic architectural protein, was first shown to bind upstream of Cp by DNA affinity pulldown, electrophorectic mobility shift assay (EMSA), DNAseI footprinting assay and chromatin immunoprecipitation (ChIP) assay (Chau 2004; Chau, 2006; Tempera, 2010; Lupey-Green, 2018). This has more recently been confirmed by ChIP-seq (Holdorf, 2011; Lupey-Green, 2018) and has led to our understanding of its importance in the regulation of Cp activity both in latency type I and type III. Initial findings suggested that CTCF was acting as a boundary element, controlling transcription from Cp (Chau, 2004). However, further investigation has shown the necessity for other CTCF binding sites to be present in the EBV genome to form chromatin loops between the OriP enhancer element and both Cp and Qp (Tempera, 2011). Hence, the multi-factorial manner by which EBV Cp transcription is controlled throughout infection is still not fully understood. We took advantage of cell lines that had previously been produced via infection with recombinant EBV in which Cp had been genetically modified (Evans, 1996) to further investigate this regulation.

Isolation of *in vivo* EBV-infected B-cells from peripheral blood usually allows their culture *in vitro* as lymphoblastoid cell lines (LCLs). Alternatively, this process can also be performed directly *in vitro* through EBV infection of isolated B-cells in the laboratory. These LCLs normally express EBV proteins consistent with latency type III and, with this in mind, the previous authors (Evans, 1996) sought to investigate the association of both the RBPJκ binding site and also glucocorticoid response elements (GREs) present between OriP and Cp. Assessment of these two sites was done independently by either deletion of the GRE region (262bp at B95-8 coordinates 10221-10482) or mutation of the RBPJκ binding site (GTGGGAAA to GTGAATTC at B95-8 coordinates 10959-10966). Deletion of the GRE region gave rise to up-regulation in Cp transcript level, whereas RBPJκ binding site mutation resulted in a modest decrease in Cp activity (Evans, 1996). Unintentionally, the deletion that removed the GREs from Cp overlaps with the now known CTCF binding site in Cp (ranging between coordinates 10401-10594 by various methods) (Chau, 2006) (Figure 1). We were therefore able to use these cell lines to investigate whether the deletion of such an intrinsic CTCF binding site upstream of Cp gave rise to changes in transcript levels associated with additional EBV promoters and whether the recruitment of transcription factors such as EBNA2 was altered in any way.

**Figure 1.**
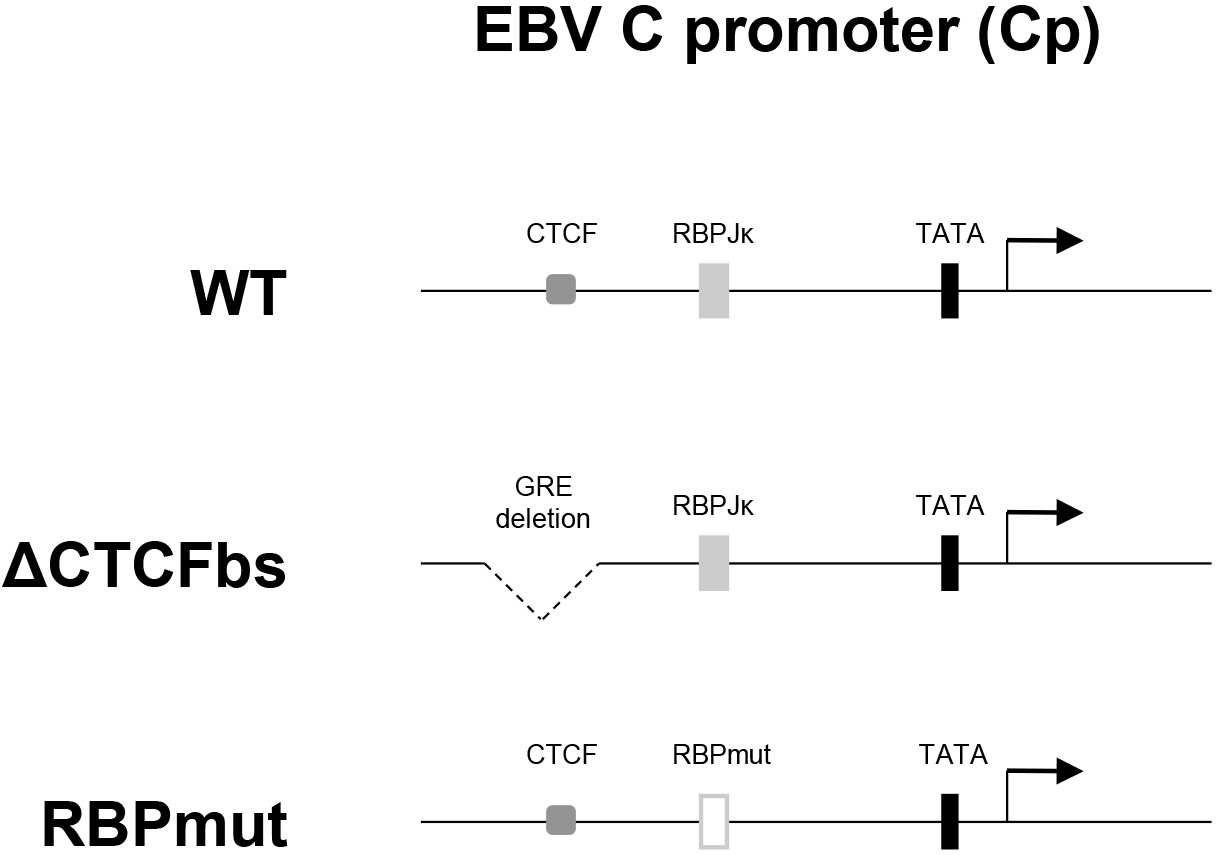
Schematic representation of modified EBV C-promoter (Cp) sequence elements in LCLs. Cell lines are described as wild type (WT), Cp-CTCF binding site deletion (ΔCTCFbs) and Cp-RBPJk binding site mutant (RBPmut) infected LCLs. ‘GRE deletion’ refers to location of the glucocorticoid response element region deletion that overlaps with the Cp-CTCF binding site.

## Results

We first tested the previous publication’s observations (Evans, 1996) of how Cp transcript levels were affected by the deletions shown in Figure 1, before assessing whether these deletions affected transcription from the alternative EBV latency-associated promoters Wp and Qp. Using qPCR we show that, relative to transcription from a wild-type B95-8 EBV infection (Figure 2A, blue bar), LCLs infected with the GRE deleted/Cp-CTCF binding site deletion (ΔCTCFbs) virus have over 5-fold greater level of transcript from Cp (Figure 2A, red bar), while only ~25% of usual Wp transcript level is present (Figure 2B, red bar). Although not statistically significant, transcription from Qp was also reduced to ~20% of wild-type level from the ΔCTCFbs virus (Figure 2C, red bar). In contrast, although the level of transcription from all promoters tested here varies somewhat from the Cp-RBPJκ binding site mutant (RBPmut) virus in LCLs (Figure 2), none of the changes are statistically significantly different than the wild-type virus level.

**Figure 2.**
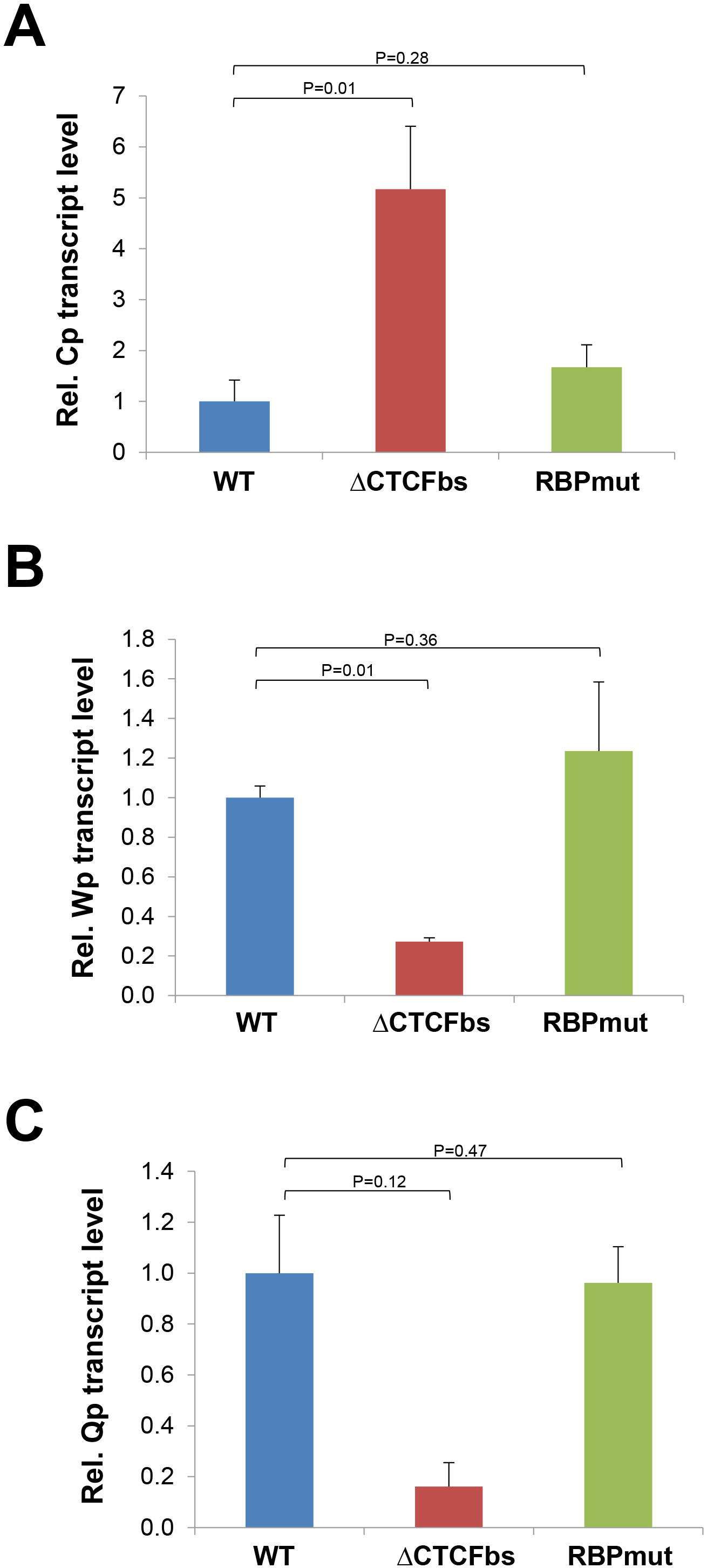
Deletion of EBV Cp-CTCF binding site leads to increased Cp transcript level in LCLs. Quantitative PCR (qPCR) analysis of FBV promoter transcript levels from (A) Cp, (B) Wp and (C) Qp, in wild-type (WT, blue bars), Cp-CTCF binding site deletion (ΔCTCFbs, red bars) and Cp-RBPJk binding site mutant (RBPmut, green bars) virus infected LCLs. Values are means (+1SD) of at least three biological replicates and two technical replicates, and relative to wild-type controls. P-value determined by Student’s T-test.

Since deletion of the CTCF binding site in Cp had caused increased transcription from Cp itself, but a decrease from Wp, we next sought to address whether this would alter protein levels of the major product EBNA2 (Figure 3). Indeed, in comparison to wild-type EBV in LCLs, EBNA2 protein level is elevated by 1.4-fold (Figure 3B, red bar). Concomitantly, expression of EBNA2 in RBPmut-infected LCLs is modestly decreased to ~80% of wild-type expression level despite both Cp and Wp showing subtle increases in transcript level (Figure 3B, green bar). This may be associated with a decrease in total protein production seen in these cell lines, as illustrated by RBPJκ expression here (Figure 3C, green bar).

**Figure 3.**
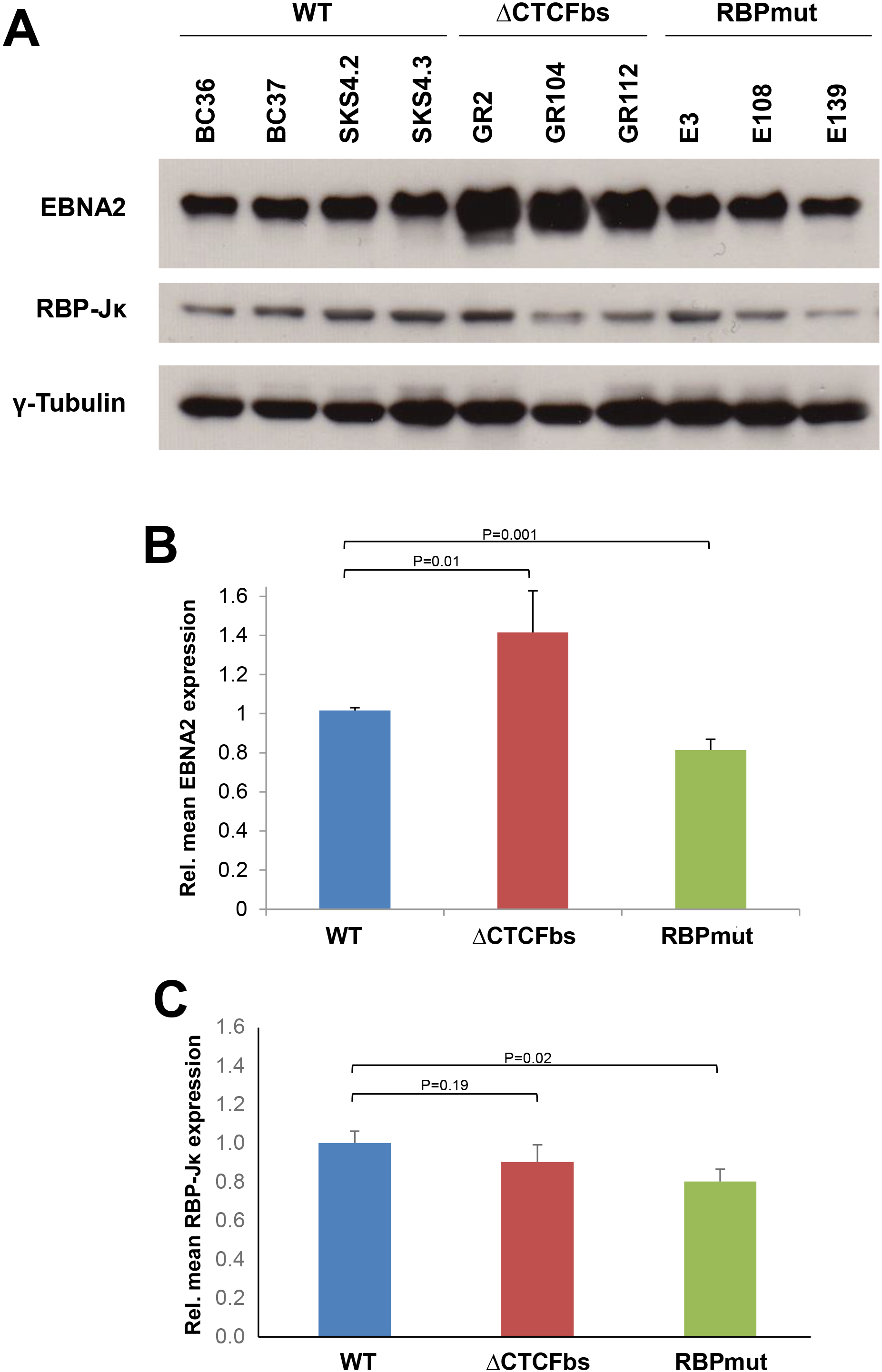
Deletion of EBV Cp-CTCF binding site leads to increased EBNA2 protein level in LCLs. (A) Western blot analysis of EBNA2 and RBPJκ protein levels, with γ-Tubulin as a loading control, in wild-type (WT), Cp-CTCF binding site deletion (ΔCTCFbs) and Cp-RBPJk binding site mutant (RBPmut) virus infected LCLs. (Representative images shown.) Semi-quantitative Image J analysis of (B) EBNA2 and (C) RBPJκ protein levels from WT (blue bars), ΔCTCFbs (red bars) and RBPmut (green bars) virus infected LCLs. Values are means (+1SD) of the biological replicates presented, from two technical replicates, and relative to wild-type controls. P-value determined by Student’s T-test.

Finally, we investigated if the level of the EBNA2 protein on the EBV genome was affected by either of the mutations of the wild-type Cp sequence (Figure 4). In comparison to the wild-type Cp (Figure 4, blue bars), deletion of the Cp-CTCF binding site results in an up to 8-fold increase in EBNA2 recruitment across Cp (Figure 4B, red bars) concurrent with a similar increase in RBPJκ recruitment (Figure 4C, red bars), while respective levels at the LMP2a promoter (LMP2ap) remain unchanged (Figures 4E-F, red bars). Interestingly, mutation of the Cp-RBPJκ binding site appears to cause a decrease in EBNA2 recruitment to Cp (Figure 4B, green bars) in comparison to wild-type infection (Figure 4B, blue bars), which is consistent with a near entire ablation of RBPJκ enrichment with the mutated promoter (Figure 4C, green bars), while association is largely unchanged at LMP2ap (Figure 4F, green bars).

**Figure 4.**
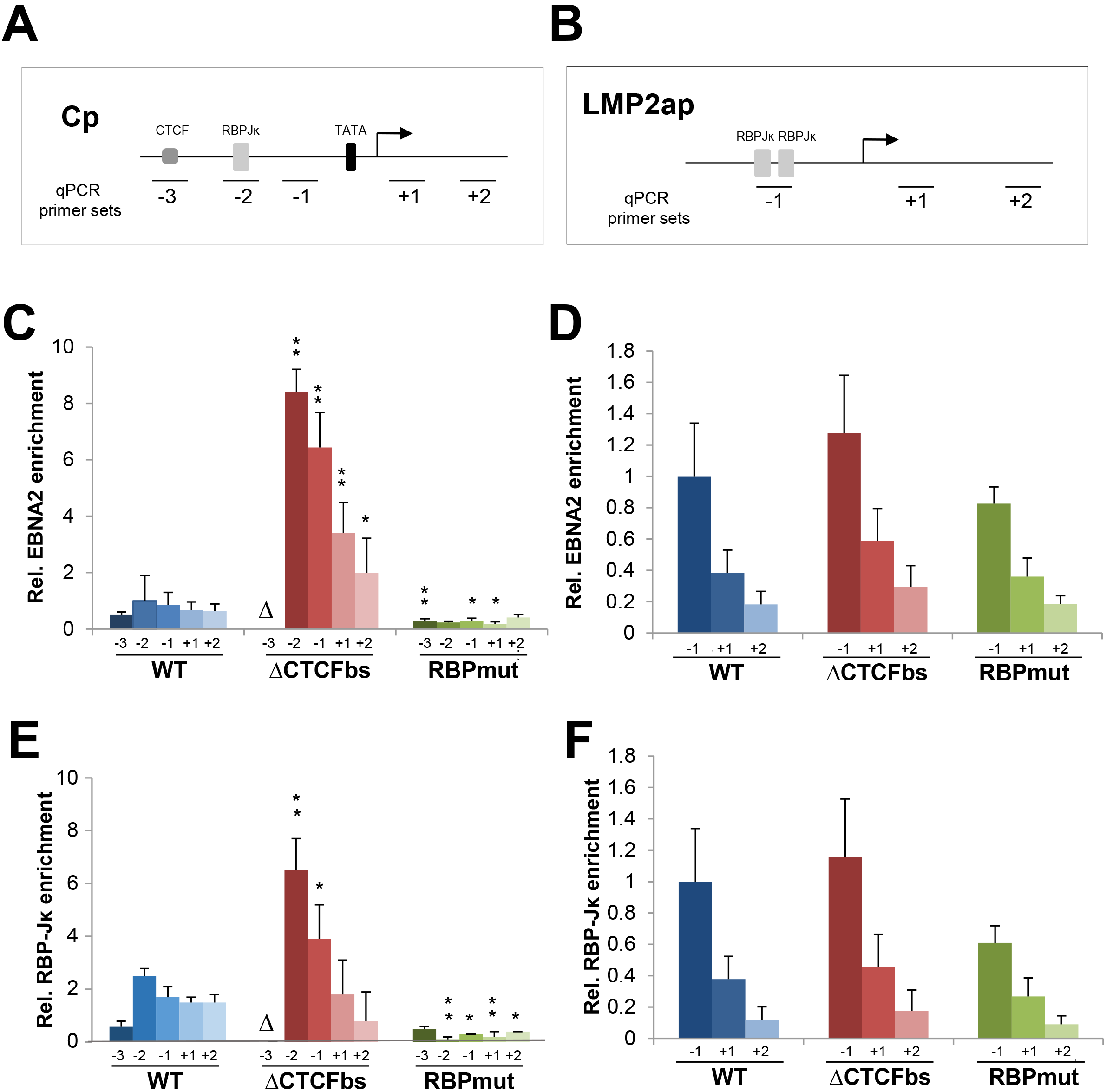
Deletion of EBV Cp-CTCF binding site leads to increased EBNA2 and RBPJκ recruitment across Cp in LCLs. Chromatin immunoprecipitation (ChIP) analysis of enrichment at (A) the C-promoter (Cp, left column) and (B) LMP2a promoter (LMP2ap, right column) of (C-D) FBNA2 and (E-F) RBPJκ in wild type (WT, blue bars), Cp-CTCF binding site deletion (ΔCTCFbs, red bars) and Cp-RBPJκ binding site mutant (RBPmut, green bars) virus infected LCLs. Schematic representations of promoters (top row) show the location of ChlP-qPCR assays relative to the transcriptional start site (angled arrow). Values are means (+1SD) of at least three biological replicates and two technical replicates, and relative to wild-type controls. P-value determined by Student’s T-test (*P<0.05; **P<0.01). Δ = deletion of this site in ΔCTCFbs LCLs.

## Discussion

The control of gene expression from the EBV genome through the regulation of transcription is known to be integral to the establishment of infection of this virus but also the shifting of latency type depending on the cellular environment that the virus finds itself in. Consequently, EBV has evolved to regulate its own expression from a small number of viral promoters that can modulate the expression level of various latency-associated genes. Through the work of a number of laboratories, we know that CTCF is an important cellular factor associated with the control of EBV gene expression. Yet, it is still incompletely understood how CTCF applies this control on promoters such as Cp. Hence, we undertook to investigate how the deletion of a CTCF binding site upstream from Cp affected transcription from various EBV promoters and also whether the association of the viral transactivator EBNA2 was in any way modulated.

We found that deletion of the CTCF binding site upstream of Cp led to up-regulation of transcription from Cp (Figure 2), as was reported in the original work with these cell lines (Evans, 1996) and for a Cp-CTCF binding site deletion mutant EBV used elsewhere (Chau, 2006). In contrast, though, the modest decrease in Cp activation that the previous authors saw with RBPJκ binding site mutation was not replicated here, where a subtle increase in transcription from both Cp and Wp was seen in RBPJmut. This difference may be due to the reduced number of cell lines available for use here, although it is important to note that our transcript data here are fully quantitative in comparison previous semi-quantitative PCRs (Evans, 1996).

Despite the decrease in the level of Wp driven transcript – which can also code for EBNA2 – associated with Cp-CTCF binding site deletion, the cumulative outcome was an increase in EBNA2 protein expression (Figure 3). As previously reported (Evans, 1996), the fold changes in transcript and protein level within cells lines do not fully correlate, adding to speculation that expression level of EBV proteins such as EBNA2 are very likely to be controlled at a post-transcriptional stage, at least in part due to the stability of both the coding transcripts and also the protein itself. The small decrease seen in EBNA2 protein level with mutation of the RBPJκ binding site in Cp, however statistically significant, appears unlikely to be functional as recruitment to the LMP2a promoter (Figure 4D, green bars) is consistent with enrichment at the wild-type promoter (Figure 4D, blue bars).

In order to investigate the mechanism by which loss of the CTCF binding site up-regulates Cp, we undertook ChIP assays to determine the association of EBNA2 with the EBV genome, as this viral transcription factor is known to transactivate Cp (Figure 4). Enrichment of EBNA2 at the wild-type Cp (Figure 4A, blue bars) was consistent to ChIP profiles across the promoter generated elsewhere (Bark-Jones, 2006) and was substantially enhanced by deletion of the CTCF binding site (Figure 4B, red bars). This increased enrichment was accompanied by an increase in RBPJκ association with Cp (Figure 4C, red bars), while mutation of the Cp-RBPJκ binding site led to modestly reduced levels of both RBPJκ and EBNA2 proteins (Figure 4B-C, green bars). This is consistent with the long-term understanding of EBNA2 recruitment to both viral and cellular genes through the RBPJκ protein and its binding sites (Ling, 1993). More recently, it has also been shown that EBNA2 itself is able to drive changes in localisation of RBPJκ, as well as the Early B-cell Factor 1 (EBF1) protein that also has a binding site within Cp (Lu, 2016). Thus, the parallel increased association of both EBNA2 and RBPJκ at the CTCF-deleted Cp is not surprising. Little change of either EBNA2 or RBPJκ association at the LMP2a promoter (LMP2ap), which itself has two RBPJκ binding sites and an EBF1 binding site, is consistent with only subtle changes to the total cellular level of those proteins in each LCL type used here and corroborates that the high enrichment at Cp-ΔCTCFbs is due to specific recruitment (Figure 2).

Although the mechanism of increased Cp activity with CTCF binding site deletion appears to involve an increased association of EBNA2 with the promoter, the full explanation for this phenomenon remains enigmatic. As EBNA2 expression can be driven from Cp, it becomes something of a ‘chicken and egg’ situation to unravel how the two are associated and whether some form of positive feedback loop exists. Despite the lack of ChIP data to confirm that the Cp deletion resulted in the loss of CTCF binding directly upstream of Cp-ΔCTCFbs, the ability of EBNA2 to associate with this location appears to be the more likely reason for concurrent up-regulation of Cp, with Cp enrichment (~8-fold) substantially greater than the rise in EBNA2 protein level (~1.4-fold - Figure 3A). Of course, we cannot preclude that deletion of the GRE region encompassing the CTCF binding site has in some way affected the association of other repressive factors between OriP and Cp, or indeed allowed the recruitment of activators other than EBNA2. It is unlikely that the loss of the GRE sites specifically has led to some form of higher Cp activation, since they appear to only have the ability to stimulate expression (Sinclair, 1994). Indeed, the finding that another Cp-CTCF binding deletion virus (Chau, 2006: coordinates 10393-10590) with only partial overlapping excised sequence to the virus used here (Evans, 1996: coordinates 10221-10482) displays the same up-regulation of Cp activity support our belief that this effect is a direct result of CTCF loss between OriP and Cp.

The ability of EBNA2 to increase association with this locus after Cp-CTCF binding site deletion may well be due to changes in the three-dimensional (3D) organisation of the EBV genome. Studies of other CTCF binding site mutants have shown that usual looping to the OriP enhancer driven by CTCF molecules can be disrupted and results in changes to EBV promoter transcription. Abrogation of Qp looping to OriP by deletion of the Qp-CTCF binding site – Qp is active in type I latency – caused up-regulation of Cp transcription, but deletion of the Cp-CTCF binding site led to a decrease in looping from oriP to both promoters (Tempera, 2011). It is tempting to hypothesise that this more accessible 3D structure may allow higher levels of recruitment of EBNA2, along with RBPJκ, to Cp under these circumstances, which subsequently leads to increased Cp transcription. Chromosome conformation capture (3C) assays would first be necessary to confirm that looping between OriP-Cp has indeed been modulated in this fashion prior to further studies. Additionally, similar analyses of a Cp-CTCF binding site deletion in an EBNA2 knockout EBV background or an EBNA2-depletion model might allow further investigation of whether higher EBNA2 recruitment is fully necessary to drive increases to Cp transcript level or whether removal of the CTCF binding site alone is enough to conserve this phenotype.

Several studies have shown that depletion of CTCF protein using short interfering RNA (siRNA) methods leads to increased Cp activity in wild-type EBV-infected 293 cells, whereas Qp transcript levels decrease (Tempera, 2011). In fact, CTCF depletion in type I latency Mutu cells caused increased EBNA2 expression, while conversely CTCF over-expression led to a decrease of EBNA2 protein in type III latency Raji cells (Chau, 2006). Indeed, greater total levels of CTCF protein are usually found in type I latency cells than in type III latency cells, supporting a model that increased binding of CTCF to the EBV genome correlates with decreases in transcription from Cp (Chau, 2006; Hughes, 2012). Therefore, it appears that CTCF may be involved in the regulation of latency type during EBV infection. It has been reported that another mutant EBV genome without the Cp-CTCF binding site was able to shift to latency type I, despite showing continued expression from Cp in comparison to the wild-type virus, in a B-cell superinfection model (Hughes, 2012). However, total cellular CTCF levels were not analysed here. Thus, it appears that, although CTCF contributes to the establishment and restriction of latency type, it may not be essential for maintenance of latency type.

Nevertheless, the data presented here support previous observations that binding of CTCF directly upstream of Cp is important for the regulation of transcription from this and other EBV promoters. Deletion of the Cp-CTCF binding site leads to a higher level of Cp transcripts and EBNA2 protein, likely through increased transactivation via higher recruitment of EBNA2, which itself becomes possible due to a more open and accessible 3D chromatin structure through the removal of the large zinc-finger protein and abrogation of normal OriP-Cp looping. Taken together, we confirm the importance of CTCF as a genomic architectural organising protein in the regulation of viral transcription and gene expression during the establishment and, to some extent, maintenance of latency type during EBV infection.

## Materials & Methods

### Cell lines

Established lymphoblastoid cell lines (LCLs) used in this study were first described elsewhere (Evans, 1996) and were a kind gift from Prof. Paul Farrell (Imperial College London). In short, wild-type EBV-infected LCLs (WT; BC36, BC37, SKS4.2, SKS4.3) acted as controls for Cp-CTCF binding site deleted EBV-infected LCLs (ΔCTCFbs; GR2, GR104, GR112) and Cp-RBPJκ binding site mutated EBV-infected LCLs (RBPmut; E3, E108, E139). All cells were grown in RPMI supplemented with 10% FCS, penicillin and streptomycin (Sigma) and were split 1:3 twice a week to maintain growth in culture.

### Quantification of EBV transcript level

RNA was extracted from approximately 1×10^6^ cells using the RNeasy mini kit (Qiagen) following the manufacturer’s instructions. For all samples, 1μg of each RNA sample was reverse-transcribed to cDNA using SuperScript III First-Strand Synthesis Supermix (Invitrogen). Around 1% of the product was then used per qPCR reaction, which was performed on an ABI 7900HT real-time PCR machine using previously published EBV promoter-specific primers/probe combinations and cellular controls (Bell, 2006) with the Taqman low ROX Probe 2X MasterMix (Eurogentec). Dissociation curve analysis was performed during each run to confirm absence of non-specific products. Data are representative of three individual experiments and averaging of biological replicates.

### SDS-PAGE and Western blotting

In short, protein extracts were resolved by sodium dodecyl sulphate-polyacrylamide gel electrophoresis (SDS-PAGE) and transferred to nitrocellulose membranes before Western blot analysis was performed using an ECL kit (Amersham) for visualization of protein levels, all as described previously (Anderton, 2007). Protein extractions were performed at least twice and representative results are shown. Primary antibodies used were: monoclonal antibodies against EBNA2 (clone PE2; DAKO), RBPJκ (ab25949; Abcam) and γ-Tubulin (T6557; Sigma). Semi-quantitative analysis of protein levels was carried out using Image J software and comparative densitometry to γ-Tubulin loading levels.

### Chromatin Immunoprecipitation

Chromatin immunoprecipitation (ChIP) assays were carried out using a ChIP Assay Kit (17–295; Millipore) according to the manufacturer’s instructions, as described previously (Paschos, 2009). Chromatin was sheared to a range of 200-800bp in length from 1×10^6^ cells per ChIP in 200 ml of lysis buffer using a Bioruptor sonicator (UCD-200; Diagenode) on a high setting for a total of 12 min (30 sec ‘on’/30 sec ‘off’ intermittent sonication). Chromatin was immunoprecipitated using either EBNA2 (ab90543; Abcam) or RBPJk (ab25949; Abcam) specific antibodies, with normal mouse or rabbit IgG serum as negative controls, respectively (12-371, 12-370; Millipore). Isolated DNA was assayed by qPCR using the Platinum SYBR Green qPCR SuperMix (11733; Invitrogen) on an ABI 7900HT realtime PCR machine. Using standard curves, 1% of input was compared to the immunoprecipitated DNA sample and the values from the IgG negative control were subtracted as background. The data are representative of two independent experiments with averaging of all biological replicates. Sequences of the primers used for ChIP-qPCR are listed in Table 1.

**Table 1.**
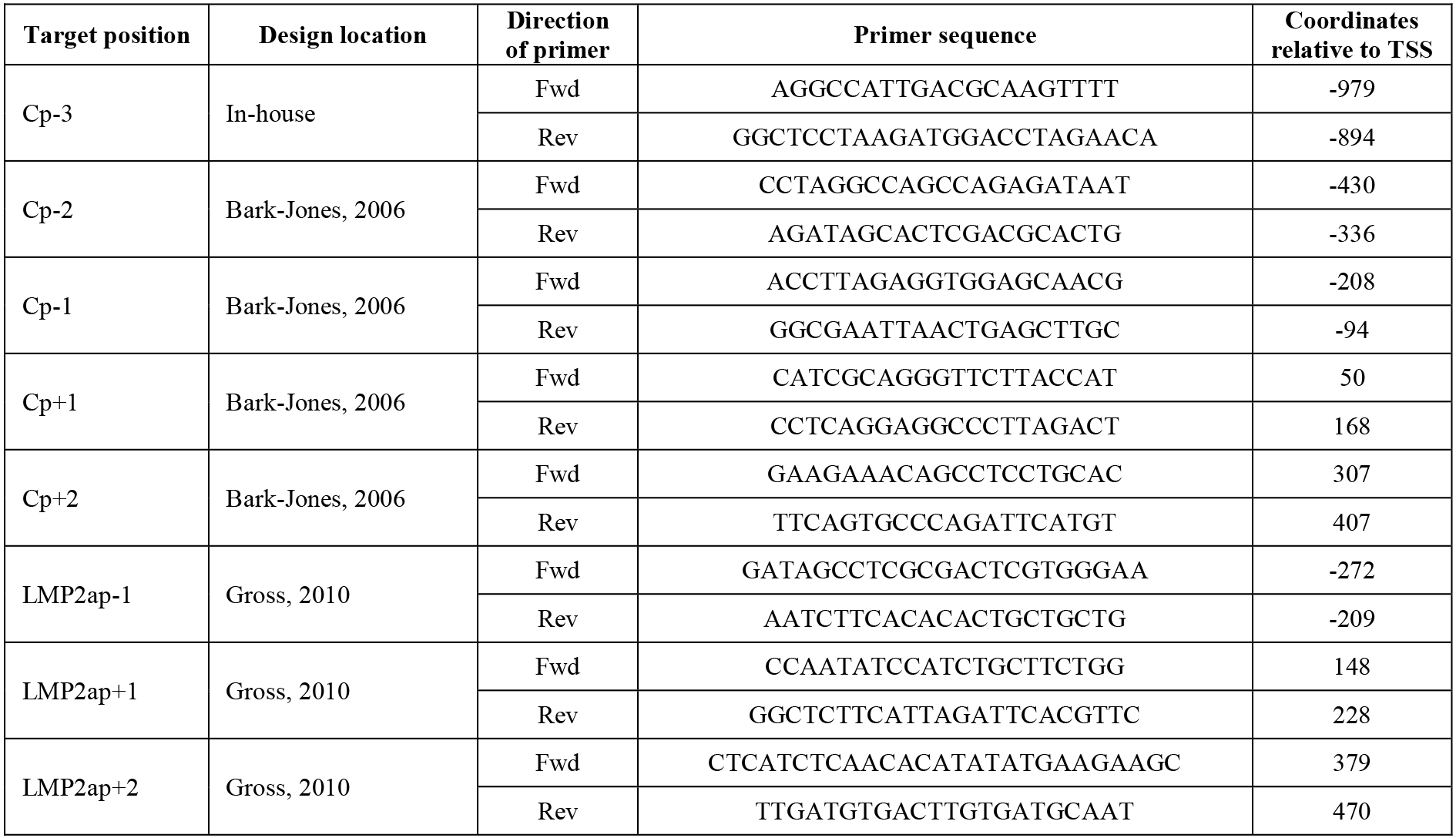
Primers used in this study for ChIP-qPCR.

## Author contributions

MJA and IJG conceived and designed the study. IJG carried out all data analysis, interpretation and wrote the manuscript.

## Acknowledgements

This manuscript was written *in memorium* of Prof. Martin J Allday and also towards the memory of our colleague and friend Dr Mark Bain. Thanks to Dr Rob E. White for helpful discussions and critical review of the manuscript. This project was funded by project grant 049293 awarded to MJA/IJG by the Wellcome Trust.

